# The effect of terminal globular domains on the response of recombinant mini-spidroins to fiber spinning triggers

**DOI:** 10.1101/2020.02.13.940478

**Authors:** William Finnigan, Aled D. Roberts, Nigel S. Scrutton, Rainer Breitling, Jonny J. Blaker, Eriko Takano

## Abstract

Spider silk spidroins consist of long repetitive protein strands, flanked by globular terminal domains. The globular domains are often omitted in recombinant spidroins, but are thought to be essential for the spiders’ natural spinning process. Mimicking this spinning process could be an essential step towards producing strong synthetic spider silk. Here we describe the production of a range of mini-spidroins with both terminal domains, and characterize their response to a number of biomimetic spinning triggers. Our results suggest that the inclusion of the terminal domains is needed to match the response to shear that native spidroins exhibit. Our results also suggest that a pH drop alone is insufficient to trigger assembly in a wet-spinning process, and must be combined with salting-out for effective fiber formation. With these insights, we applied these assembly triggers for relatively biomimetic wet spinning. This work adds to the foundation of literature for developing improved biomimetic spinning techniques, which ought to result in synthetic silk that more closely approximates the unique properties of native spider silk.

## 1. Introduction

Spider dragline silk has impressive mechanical properties, with high strength and good extensibility resulting in a level of toughness, which exceeds all other natural and synthetic fibers ^1^. However, unlike silk worms, spiders cannot be efficiently farmed for their silk ^2^. For this reason, the production of recombinant spider silk proteins (spidroins), and their subsequent spinning into synthetic spider silk fibers, has been an active topic of research for a number of decades ^3^.

Major ampullate spider silk proteins (spidroins) are typically 200–350 kDa in size and constitute the dragline silk of spiders. Generally, spidroins have three distinct regions (**Figure 1**) ^3^. The vast majority of the protein is repetitive, consisting of alternating polyalanine regions and glycine-rich regions ^4^. At the terminals of the spidroin exist non-repetitive domains, referred to as the N- and C-terminal domains (NTD and CTD). These globular terminal domains are crucial in facilitating the soluble storage of the spidroins at high concentrations (30 – 50 % w/v) in the silk gland, and in initiating fiber assembly ^5^.

**Figure 1.**
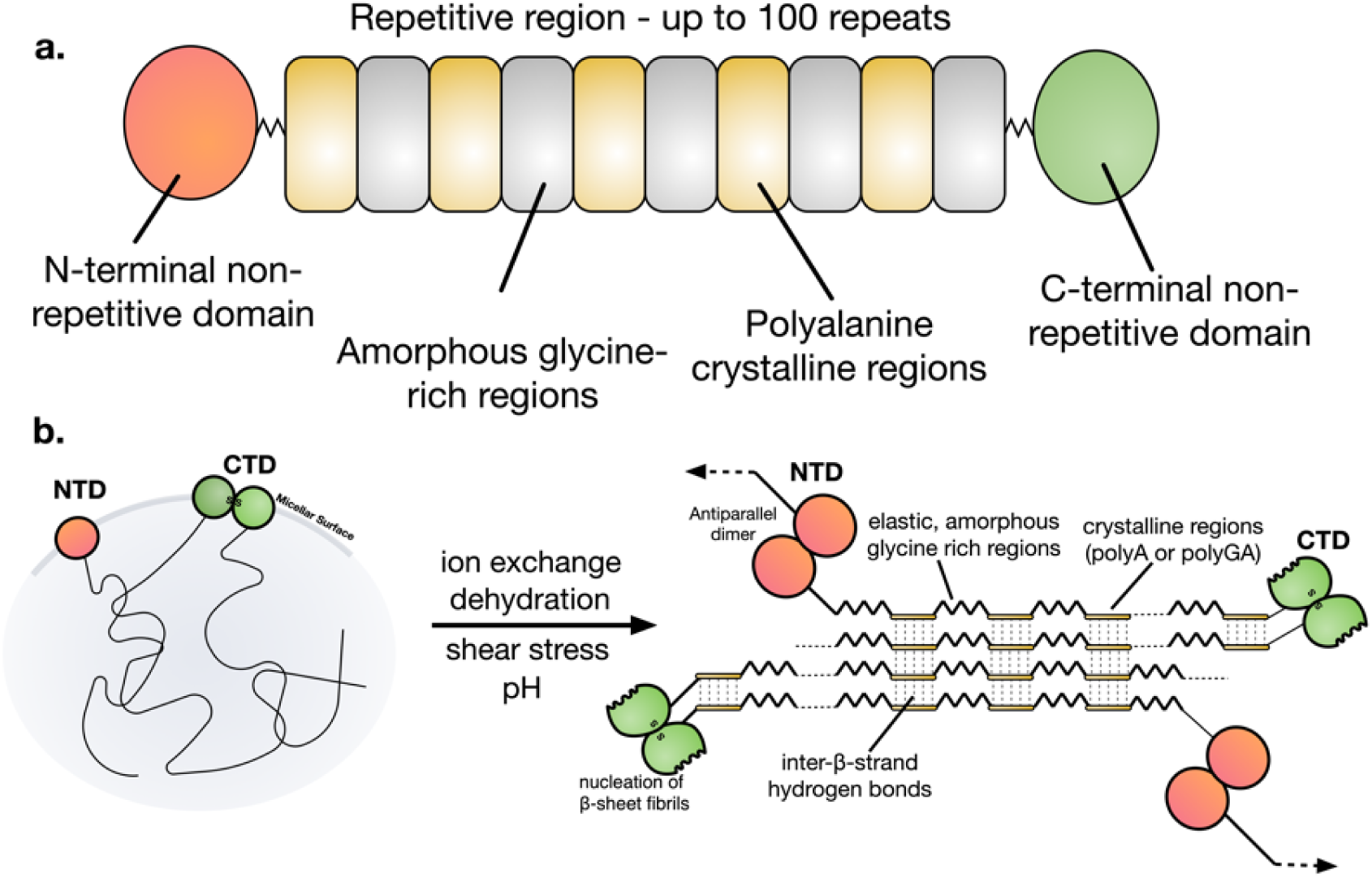
Schematic representation of a native spider silk protein. **A.** The domain structure of spider silk proteins, consisting of non-repetitive N- and C- terminal domains, flanking a much larger repetitive section, which alternates between glycine-rich regions and polyalanines. **B**. A model for the conversion of soluble spidroins, stored as protein micelles, into insoluble silk fibers through the assembly triggers of shearing force or elongational flow, changing pH, dehydration and changing salts in the silk gland of a spider. Reproduced from ^45^.

Much of the research into recombinant spider silk has focused on spidroins consisting of only the repetitive region. Whilst some of the largest recombinant spidroins have resulted in fibers with good mechanical properties, larger repetitive regions typically result in poor spidroin yields ^6,7^. Commonly, denaturing conditions have been employed in either the purification or spinning processes. Silk proteins purified under these conditions have been shown to lack the response to shear native spinning dopes exhibit, essential in the use of shear to provide alignment in the fiber, and as an assembly trigger itself ^8^. In contrast, biomimetic spinning utilising correctly folded terminal domains may offer the production of more biomimetic spinning dopes, and a route to synthetic spider silk fibers with superior mechanical properties ^3,5^.

In spiders, dragline spidroins are stored in an ampulla (or sac) in the silk gland, where the highly concentrated spinning dope forms a lyotropic liquid crystalline solution ^9,10^. Upon spinning, the spidroins proceed through a long and increasingly narrow S-shaped spinning duct where the coordinated action of acidification, ion exchange, dehydration, shearing force and elongational flow, is proposed to trigger assembly and promote alignment of β-sheet nanocrystals as the fibers are formed (**Figure 1**) ^10,11^. Chaotropic sodium and chloride ions are replaced with potassium and kosmotropic phosphate ions during the spinning process, inducing salting out of the spidroins ^12–14^. Chaotropic ions have been shown to prevent intra- and intermolecular interactions on the recombinant repetitive regions, while kosmotropic ions promote hydrogen bond interactions in the glycine-rich regions ^15^. The pH drops from pH 7.6 at the beginning of the duct, to pH 5.7 by halfway through, and likely lower near the spinneret, as a result of the action of a carbonic anhydrase ^16,17^. The decreasing pH causes conformational changes in the N- and C-terminal domains, which act as regulatory elements for the control of spidroin assembly ^3^. In contrast, the molecular structure of some recombinant repetitive regions have been shown not to respond to pH ^18^.

The NTD is known as the ‘lock’, as this domain dimerises in response to the decreasing pH ^19^. This dimerization ‘locks’ the spidroins into an infinite network, as the CTDs form a disulphide-linked dimer ^20–22^. The CTD is proposed to partially unfold in response to decreasing pH. This change is thought to cause the CTD to form β-sheet amyloid fibrils, nucleating the formation of β-sheet fibrils in the repetitive region, in a process analogous to the nucleation of various kinds of amyloid fibers ^20,21^.

Recent work showed a small mini-spidroin featuring both terminal domains could be spun into a fiber by wet-spinning using a coagulation bath at pH 5.0, rather than the more commonly used denaturing methanol or isopropanol ^5^. Here we build upon this to further investigate the expression of a range of mini-spidroins featuring pH-responsive terminal domains, which we have termed “complete” mini-spidroins to differentiate them from mini-spidroins consisting of only a repetitive region. We demonstrate the effects of pH, ion exchange and shearing force, which spiders employ during spinning, on one of these complete mini-spidroins and identify potentially relevant triggers for the development of better biomimetic spinning techniques. Finally, we use the resulting understanding of these assembly triggers to biomimetically spin synthetic spider silk fibers.

## 2. Methods

### 2.1 Spidroin cloning, protein expression and purification

Genes for the N- and C-terminal domains (NTD1, NTD2, CTD1 and CTD2) were synthesised and cloned into the *Nco*I and *Hind*III restriction sites of the plasmid pNIC28-BSA4 ^23^. The plasmid designated pTE1253 (NTD2-BsaI-CTD1-pNIC28) was generated by Gibson assembly ^24^. R_18_ was gene synthesised with the inclusion of *Bsa*I and *Bpi*I restriction sites. Repetitive regions below R_18_ in size were generated by PCR with the inclusion of *Bsa*I and *Bpi*I restriction sites, and cloned directly into pTE1253 (Supplementary Material **Figure S4**). Repetitive regions larger than R_18_ were generated according the scheme outlined in Supplementary Material **Figure S3**, before sub cloning into pTE1253. All constructs featured an N-terminal 6x His tag. Plasmid sequences are available as supplementary material (**Supplementary Material 3 zip**). Protein sequences are available at the end of the Supplementary Material as Supplementary Table 2. Cloning was carried out in *E. coli* 5α.

Protein expression was carried out in *E. coli* BL21(DE3) in Terrific Broth media with the addition of 100 μg/μl kanamycin. Cells were grown to approximately 0.8 OD_600nm_ (optical density at 600 nm) at 37 °C with shaking at 180 rpm, at which point IPTG was added to a concentration of 200 μM. Temperature was dropped to 20 °C for protein expression overnight. Cell lysate was prepared by sonication on ice followed by centrifugation to remove the insoluble fraction. Proteins were purified from cell lysate by immobilized metal affinity chromatography using a Ni-NTA resin, eluted using 250 mM imidazole. Purified protein was dialyzed twice against 25 mM TrisHCl pH 8.0, at 4 °C. Aggregated protein following dialysis was removed by centrifugation. Protein expression and purification was analyzed by SDS-PAGE. Protein concentrations were determined in triplicate by OD_280nm_ using a Nanodrop 2000 (Thermo Scientific), using an extinction coefficient and molecular weight for each protein calculated using the ExPaSy ProtParam tool ^25^. Where necessary dilutions of proteins were made before determining the concentrations. Single use aliquots of protein were stored at −80 °C where appropriate.

### 2.2 Size exclusion chromatography of NTD2

Buffers consisting of 25 mM Tris pH 8.0 with either 0 or 300 mM NaCl, and 25 mM MES pH 5.5 with either 0 or 300 mM NaCl, were prepared and filtered. 250 μl of purified NTD2 at 3.4 mg/mL was loaded onto a Superdex 200 Increase 10/300 GL size exclusion column pre-equilibrated in the relevant buffer, using an AKTA Pure system. The sample was eluted over 1.2 column volumes, with samples coming off the column monitored at OD_280nm_. Retention times were compared to a standard curve of known proteins, and the theoretical molecular weight of NTD2 to estimate oligomeric state.

### 2.3 Tryptophan fluorescence

Tryptophan fluorescence for NTD2 at 0.8 mg/mL was recorded between 310 and 400 nm (bandwidth 10 nm) while exciting at 280 nm (bandwidth 20 nm). The assay was performed in triplicate in a 96-well microtiter plate using a M200 Infinite plate reader (Tecan). pH values between 8.2 and 5.5 were achieved using assay concentrations of 50 mM HEPES, Tris-HCl, MES or sodium acetate, depending on the desired pH. Blank measurements were recorded in every condition and subtracted from the final reading.

### 2.4 Dynamic scanning fluorimetry

Dynamic scanning fluorimetry assays were carried out using 96-well PCR plates (BioRad) in a CFX Connect Real-Time PCR detection system (BioRad), using the HEX filter ^26–28^. Assay volume was set at 30 μL Suitable protein concentrations between 1 – 10 μM for each assay were determined by titrating the amount of protein. A master mix of protein and sypro orange was prepared, and 5 μL added to every well. Sypro orange was used at an assay concentration of 5x. 25 μL of each assay buffer solutions were transferred by multi-channel pipette to the assay plate in triplicate. The assay plate was sealed and briefly centrifuged before starting the assay. After a 3 minute hold at 25 °C, temperature was increased by 0.4 °C every 30 seconds up to 95 °C. Peaks in dF/dt, corresponding to the T_M_ of the protein, were identified using the Biorad CFX Manager software.

To investigate the response of CTD1 to pH, a range of pH values between 8.2 and 5.5 were set up in a 96 deep-well block to achieve assay concentrations of 50 mM HEPES, Tris-HCl, MES or sodium acetate. No salt, or assay concentrations of 250 mM NaCl or 250 mM KCl were added to three separate sets of buffers. Subsequent assays used assay concentrations of 40 mM HEPES (pH 8.0 and 7.0), 40 mM MES (pH 6.0), and 40 mM sodium acetate (pH 5.0). NaCl or KPi concentrations were added at each pH for assay concentrations between 0 and 400 mM. Potassium phosphate was prepared at each pH from solutions of KH_2_PO_4_ and K_2_HPO_4_, referred to as KPi here.

### 2.5 Rheology flow sweeps

Rheology was conducted using 180 μL sample of 200 mg/mL the mini-spidroin designated N-R_7_-C for shear sweep measurements, or 100 mg/mL N-R_7_-C for the frequency, amplitude and time sweeps. The lower concentration of 100 mg/mL was used initially to facilitate more preliminary experiments. A discovery HR-2 hybrid rheometer (TA Instruments) was used, with a parallel plate geometry with a plate diameter of 20 mm. A geometry gap of 500 μm was used and a solvent trap attached to prevent evaporation. Experiments were carried out at 25 °C. The viscosities of the solutions under shear sweeps were investigated using a logarithmic steady shear rate increase from 0.01 to 1000 s^-1^. Samples were run four times each, with and without a 15 min settle time to check for the effect of immediate consecutive runs.

### 2.6 Turbidity assays to investigate aggregation

An array of buffer conditions were prepared in a 96-well microtitre plate. Assay concentrations of 50 mM HEPES (pH 8.0 and 7.0), 50 mM MES (pH 6.0), or 50 mM sodium acetate (pH 5.0) were prepared, each with 0 to 250 mM assay concentrations of NaCl or KPi, in triplicate. KPi stock solutions were prepared at each pH from KH_2_PO_4_ and K_2_HPO_4_. Assays were initiated by the addition of 25 μL protein for an assay concentration of approximately 40 μM, gently mixed by tapping the plate before placing into the Clariostar plate reader (BMG Labtech). To measure changes in turbidity, the average of four OD_340_ readings was recorded every minute per well for one hour. Protein concentrations before and after the assay were determined by taking 2 μL of protein from the top of each well and measuring OD_280_ using a nanodrop.

### 2.7 Analysis of mini-spidroin assembly

To investigate the effect of potassium phosphate during a wet spinning process, buffers containing 50 mM HEPES pH 8.0, or 50 mM sodium acetate pH 5.0 were prepared with the addition of 0 to 500 mM potassium phosphate, prepared from KH_2_PO_4_ and K_2_HPO_4_ at the relevant pH. N-R_7_-C at 100 mg/mL was extruded at 0.5 mL/hr using a syringe pump (Cole-Palmer 74900 series) with a 1 mL syringe, through a blunted 16G needle.

### 2.8 Fiber spinning and analysis

N-R_7_-C at 300 mg/mL was extruded using a syringe pump through a pre-pulled glass capillary (MGM-3-1.5-5NF, 30 μm tip, FivePhoton Biochemicals) at 0.5 mL/h using a syringe pump (Cole-Palmer 74900 series) with a 1 mL syringe into a coagulation bath of 500 mM sodium acetate pH 5.0, 200 mM NaCl. A fiber was pulled using tweezers onto a custom made rotating collector onto which fibers were collected continuously at approximately 9.6 m/min.

Individual fibers were mounted onto cardboard mechanical testing windows with a 5 mm gauge length using scotch tape. Each fiber on a window was imaged by light microscopy (Leica DMI6000) at 20x magnification. Fiber diameters were measured using ImageJ at multiple points along the fiber, and the average taken. The cross sectional area of each fiber was calculated from the average diameter, for subsequent mechanical testing measurements. Fibers on cardboard frames were mounted into a tensile testing machine (Instron 3344; Instron Ltd.), equipped with a 10 N load cell. Upon mounting, the sides of the window were cut and the fiber loaded. Tensile tests were performed at a rate of 0.5 mm/min at room temperature and humidity (28 °C, 52 % humidity). Engineering stress was calculated from the measured load using the calculated cross-sectional area. The ultimate tensile stress (UTS) and strain to failure were determined, Young’s modulus was calculated from initial linear portion of the stress-strain curve, and toughness calculated from the area under the stress-strain curve.

Fiber diameters and morphologies were assessed using scanning electron microscopy (SEM) for spidroins spun into a coagulation bath consisting of 500 mM sodium acetate pH 5.0, 200 mM NaCl. A number of these fibers were mounted onto aluminium SEM studs with double-sided conductive carbon tape and sputter-coated with gold/palladium (Gatan Model 682 Precision Etching Coating System, USA). Fibers were imaged using scanning electron microscopy (SEM) on a Hitachi S300 N SEM and an FEI Quanta 250 FEG-SEM. Samples were initially sputter coated (10 nm thickness) with an Au/Pd alloy to enhance conductivity.

Fiber crystallinity was determined by wide-angle X-ray diffraction (WAXD) using a PANalytical X’Pert Pro (UK) instrument with Kα radiation source (Kαav = 1.542 Å). The fiber bundle was mechanically attached to a zero-background holder and the diffraction angle ranged between 5 – 60° with a scanning rate of 1° min^-1^.

## 3. Results and Discussion

### 3.1 Functional terminal domains are necessary to produce complete mini-spidroins for biomimetic spinning

We first sought to identify and characterize highly expressed, soluble and pH-responsive N- and C-terminal domains, which we selected from major ampullate spidroins 1 and 2 of *Latrodectus hesperus* (Supplementary Material **Figure S1**, NTD1, NTD2, CTD1 and CTD2 for major ampullate spidroins 1 and 2 respectively. NTD2 and CTD1 were used for this study). A tryptophan residue, which is buried in the monomer of NTDs, has previously been shown to become exposed in the dimer conformation allowing the conformational change leading to dimerization to be followed by tryptophan fluorescence ^29^. NTD2 showed a large shift in fluorescence with decreasing pH suggesting such a conformational change (**Figure 2B**). Size exclusion chromatography (SEC) showed NTD2 to form a dimer at pH 5.0, while remaining a monomer at pH 8.0, in the presence of 300 mM NaCl (**Figure 2A**). However, we noted that NTD2 eluted slightly later than expected, likely due to the addition of an unstructured section of repetitive region in the protein used in this experiment (Supplementary Material **Figure S1A**), other experiments were carried out NTD2 without this domain. In the absence of NaCl, NTD2 eluted earlier from the SEC column. In addition, following the SEC in these conditions the samples appeared visibly cloudy.

**Figure 2.**
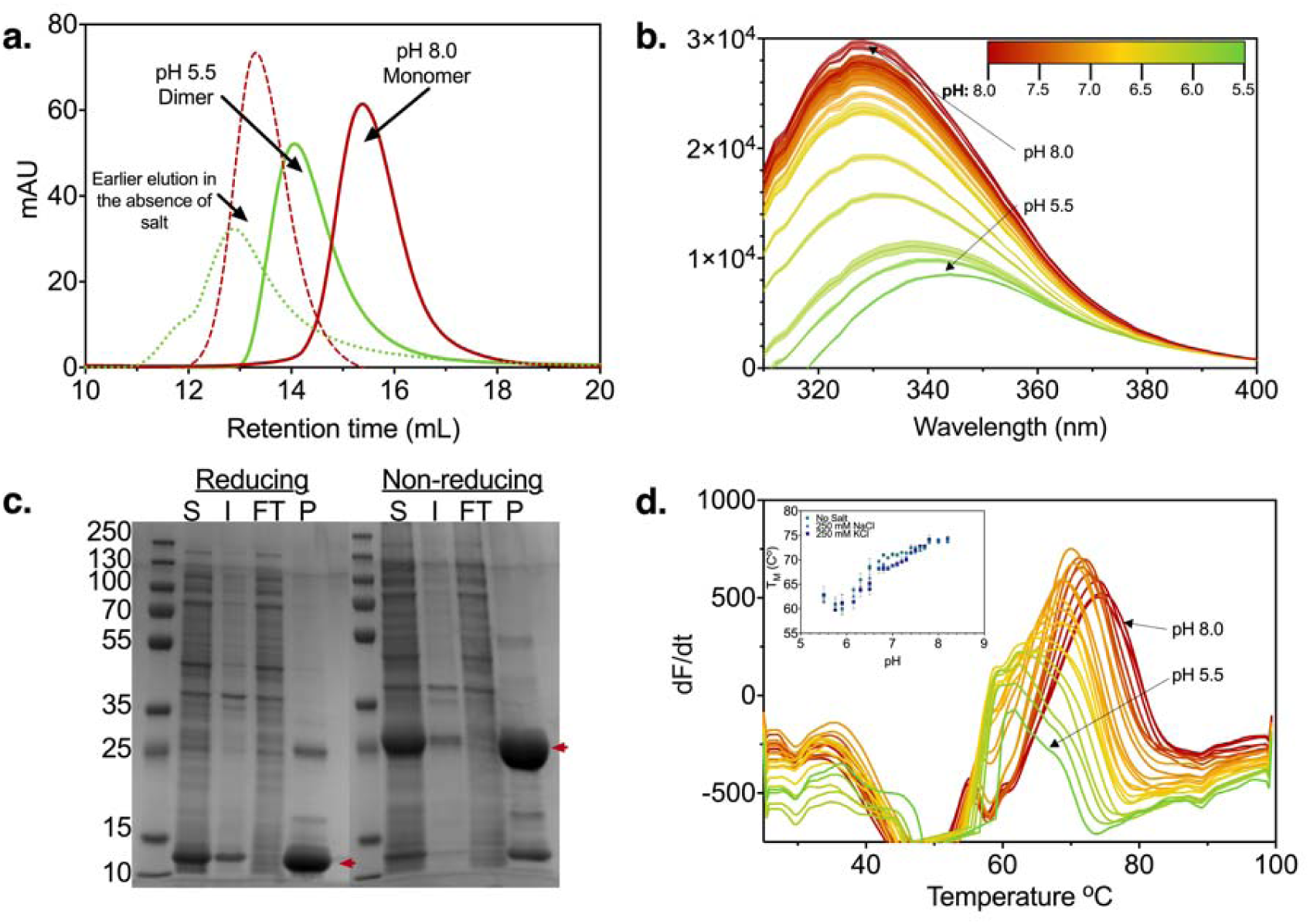
Characterisation of N- and C-terminal domains. **A**. Size exclusion chromatography of NTD2 at pH 8.0 (red) and pH 5.5 (green), showing peaks corresponding to a monomer and a dimer respectively. 250 μl of purified NTD2 at 3.4 mg/mL loaded onto a Superdex 200 Increase 10/300 GL size exclusion column pre-equilibrated in the relevant buffer. Peaks coming off the column at different retention times, corresponding to a dimer and a monomer, were detected for buffers at pH 5.5 and pH 8.0, respectively. Runs were performed in 300 mM NaCl in both cases, without which the elution time was shorter, possibly indicating larger multimers (dashed lines). Following elution, samples in the absence of salt appeared cloudy, suggesting aggregation. **B.** Tryptophan fluorescence of NTD2. **C.** Reducing and non-reducing SDS-PAGE gels showing the purification of CTD1 by nickel immobilized metal affinity chromatography. S: Soluble, I: Insoluble, FT: Flow-through, P: Purified. A band corresponding to a CTD1 monomer is observed under reducing conditions, while a band corresponding to a CTD1 dimer is observed under non-reducing conditions (red arrows). **D.** Differential scanning fluorimetry of CTD1 at various pH values pH 8.2 (red) to pH 5.5 (green), as shown in legend for B), with 250 mM NaCl. The plot shows the derivative of the fluorescence signal, with the peak corresponding to the denaturing temperature (TM) of the protein. (The assay was also performed with 250 mM KCl and without salt, giving similar results.)

CTD1 was confirmed to form a disulphide-linked dimer by non-reducing SDS-PAGE (**Figure 2C**). A spidroin featuring both NTD2 and CTD1 is therefore expected to polymerise via end-to-end linking of the terminal domains, upon dimerization of NTD2. CTD1 was shown to become less thermostable with decreasing pH by dynamic scanning fluorimetry (**Figure 2D**) ^26–28^. Higher initial fluorescence was also observed in this assay with decreasing pH (Supplementary Material **Figure S2**), indicating more exposed hydrophobic regions at lower pH values. These results correspond with CTD1 partially unfolding with decreasing pH, suggesting a sequence prone to hydrophobic β-aggregation present in CTD1 and other C-terminal domains ^20,21^, consistent with the amyloid nucleation concept described above.

Having characterized suitable terminal domains, these were incorporated into a mini-spidroin expression vector - pTE1253, into which different repetitive regions could be easily cloned for the production of complete mini-spidroins. We adopted a cloning scheme utilising Type IIS restriction sites, allowing both pseudo-scarless duplication of repetitive regions, and their transfer into pTE1253 (Supplementary Material **Figure S3** and **S4**).

### 3.2 Smaller complete mini-spidroins offer substantially higher protein yields

A range of different sized repetitive regions were generated from a codon-optimised DNA sequence for a section of the repetitive region of the major ampullate spidroin 1 from *Latrodectus. hesperus.* Repetitive regions were cloned into pTE1253 to generate a range of “complete” mini-spidroins with both terminal domains (NTD2 and CTD1) (**Figure 3A, 3B**). In addition, we generated a construct featuring both terminal domains but no repetitive region, designated NC throughout. Expression of these constructs in *E. coli* BL21(DE3) resulted in good levels of expression for constructs up to N-R_18_-C, at four hours post induction (**Figure 3C**) compared to 20h of incubation (**Figure 3D**). Expression of constructs larger than this (N-R_36_-C to N-R_291_-C) were not detected by SDS-PAGE (Supplementary Material **Figure S5**).

**Figure 3.**
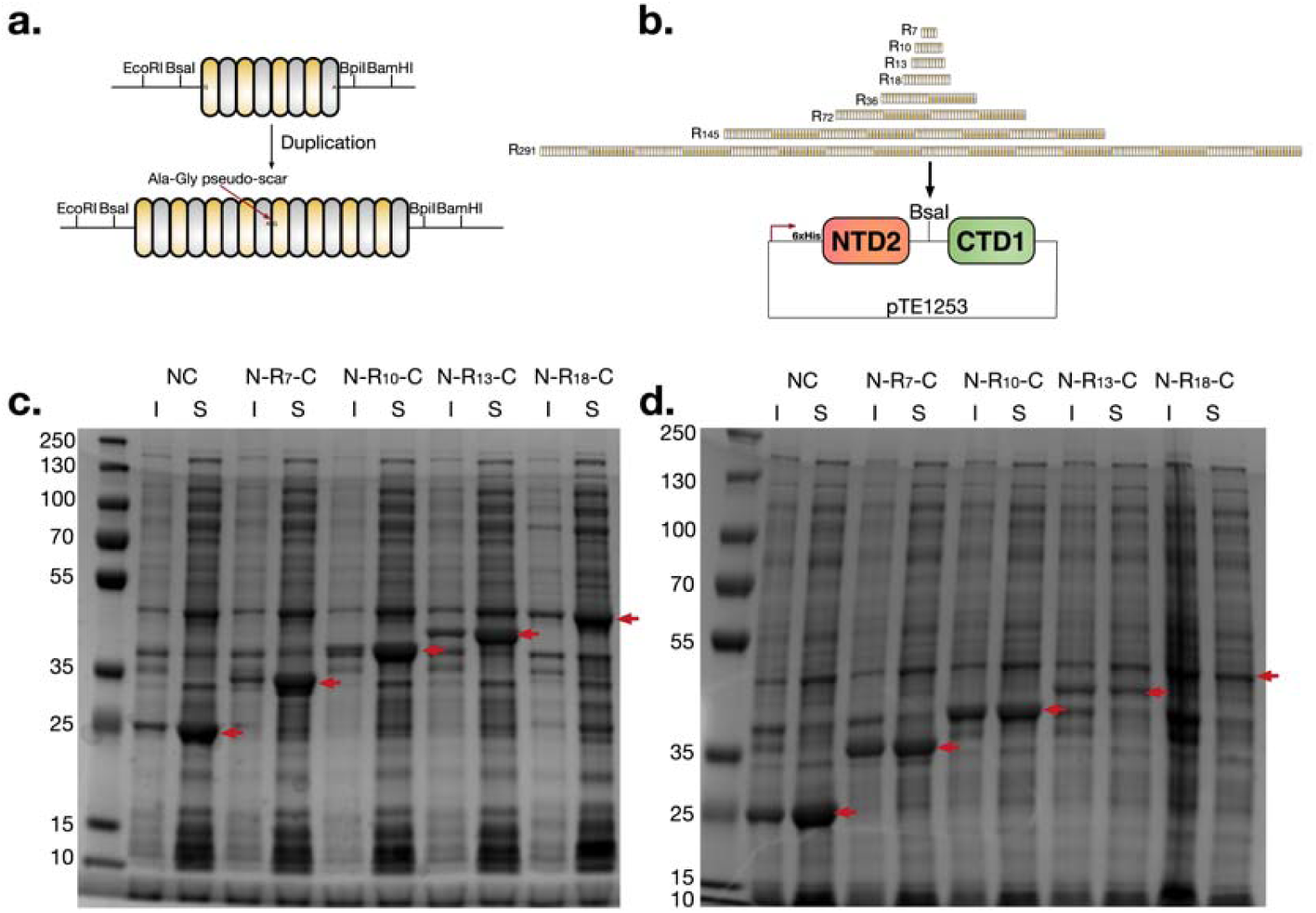
Heterologous expression of spidroins. Cloning scheme for (**A**) the duplication of repetitive regions, and (**B**) their incorporation into the pTE1253 vector for expression of complete mini-spidroins with both terminal domains. Both the duplication of repetitive regions, and their transfer into pTE1253 utilise type IIS restriction enzymes BsaI and BpiI resulting in an innocuous scar sequence which codes for alanine-glycine (Supplementary Material **Figure S4**). **C** and **D**. SDS-PAGE of soluble (S) and insoluble (I) fractions of *E. coli* lysate following expression of various mini-spidroins for four hours (C) or 20 hours (D) at 20 °C. Overexpressed proteins at the expected molecular weight are indicted by red arrows.

Smaller mini-spidroins allowed substantially higher levels of expressed soluble protein when cultures were grown overnight. In comparison, the expression level for the larger mini-spidroins decreased with longer growth times, likely due to intracellular aggregation (Supplementary Material **Figure S5**). Indeed, mini-spidroins with larger repetitive regions are proposed to be more aggregation-prone ^20^. Efforts to express the repetitive regions alone yielded no visible expression by SDS-PAGE analysis of *E. coli* lysates (data not shown).

Higher yields were obtained for the smaller mini-spidroins (∼30 mg/L purified protein for N-R_18_-C, ∼420 mg/L purified protein for N-R_7_-C), and all could be purified to high purity using nickel immobilized metal affinity chromatography (Supplementary Material **Figure S6**). Following dialysis, small mini-spidrions N-R_7_-C and N-R_10_-C could be concentrated to at least 30 % w/v as is seen in spiders ^30^, without premature aggregation using centrifugal concentrators. During this process a significantly more viscous phase was observed to form at the bottom of the concentrator. In contrast, processing of the larger mini-spidroin N-R_18_-C was not possible in this way due to aggregation. We also attempted to concentrate N-R_18_-C by reverse osmosis without success.

### 3.3 A complete mini-spidroin featuring both terminal domains displays shear thinning, similar to native spider silk spinning dopes

The rheological properties of N-R_7_-C at 200 mg/mL (20 % w/v) at pH 8.0 were investigated (**Figure 4**, Supplementary Material **Figure S7**). The spinning dope behaved as a non-Newtonian fluid, displaying shear thinning. This response to shear is shared with native spider silk and silk worm spinning dopes ^31^, but not with recombinant spidroins consisting of only a repetitive region ^32^, or with reconstituted silk fibroin (RSF) in which the terminal domains are unlikely to be correctly folded ^8,33^. This suggests that the inclusion of terminal domains is an important factor for a response to shear similar to native spider silk spinning dopes. Likely denaturing these terminal domains, as occurs in the production of RSF, might limit this response ^31,33^.

**Figure 4.**
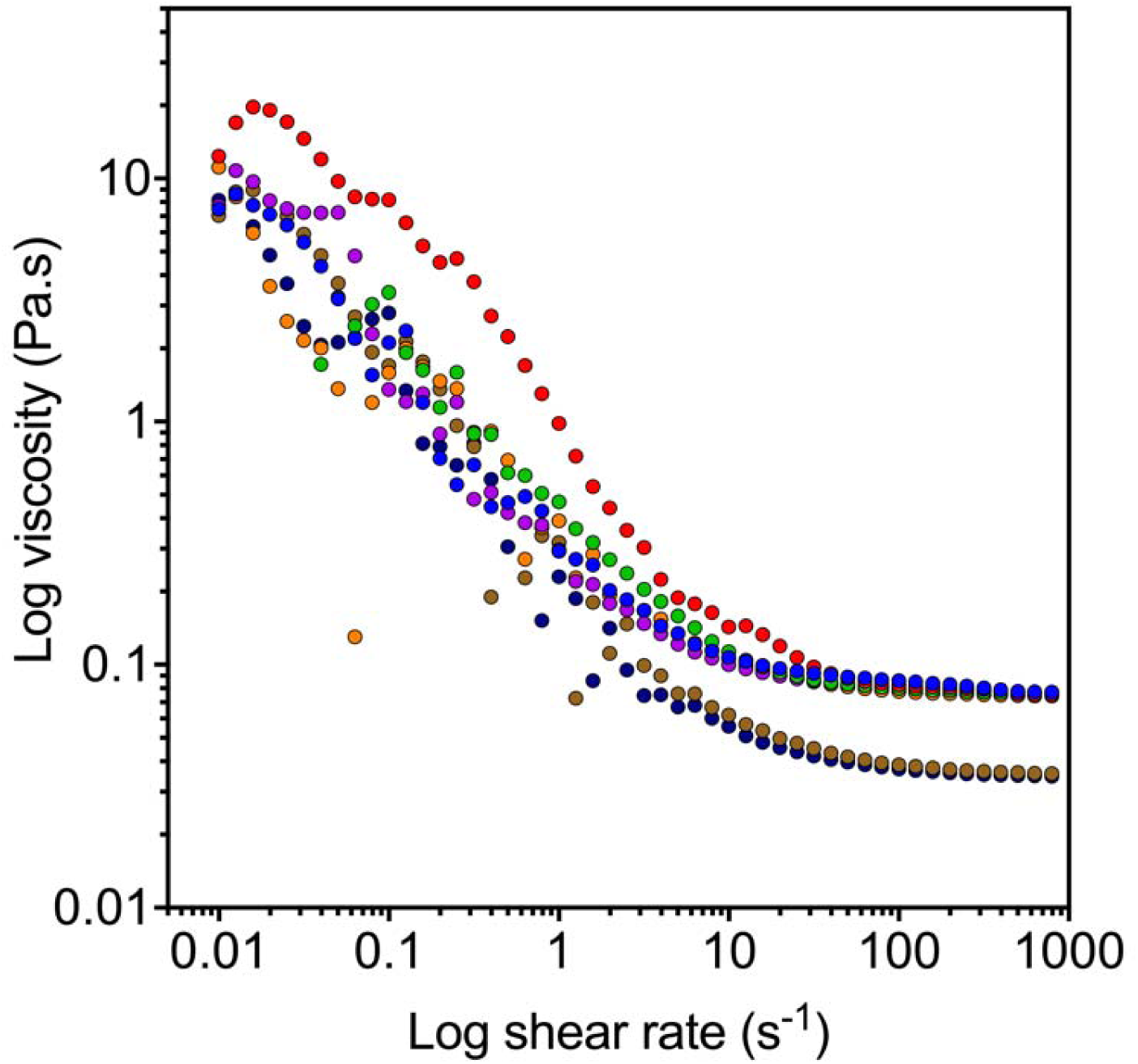
Multiple flow sweeps of 200 mg/mL (20 % w/v) N-R_7_-C. Each sample displayed shear thinning to near identical viscosity following repeated runs. Four repeated runs were carried out for two samples. Shear history did not appear to have an effect on the rheology of the sample.

At high shear rates the viscosity of our mini-spidroin plateaued, and the solution behaved as a Newtonian fluid, suggesting complete alignment of the spidroins (**Figure 4**). Low concentration native spider silk dopes show this effect, but at higher concentrations shear thickening events, which indicate shear-induced aggregation, are observed ^34^. Our mini-spidroin showed no shear-thickening events, and multiple repeated runs did not result in a difference in rheological behaviour (**Figure 4**). Taken together, these results suggest shear as an important process in providing alignment during the biomimetic spinning of complete mini-spidroins, as is thought to occur in spider silk gland ducts ^10^. However, unlike native spider silk spinning, shear does not act as an assembly trigger itself for our small mini-spidroin. This property likely facilitates their high expression levels (**Figure 3C**), and their processing to a suitably high concentration as soluble protein.

### 3.4 NTD2 requires stabilisation by electrolytes at pH 5.0 to prevent undesirable aggregation of mini-spidroins

The effect of varying concentrations of NaCl and potassium phosphate on N-R_7_-C and the terminal domains, across different pH values, was assayed by measuring turbidity and soluble protein concentration over time (**Figure 5**). Potassium phosphate was prepared at each pH from solutions of KH_2_PO_4_ and K_2_HPO_4_, referred to as KPi here. Potassium phosphate was chosen to investigate the effect of these ions, which are proposed to increase during the spinning process ^12–14^. Where aggregation occurred, an initial fast increase in turbidity was observed, followed by decreasing turbidity due to a loss of light scattering as larger aggregates formed, as reported with natural spider silk dopes ^34^. In these cases, a large decrease in the soluble protein was observed at the end of the assay (**Figure 5**, symbols).

**Figure 5.**
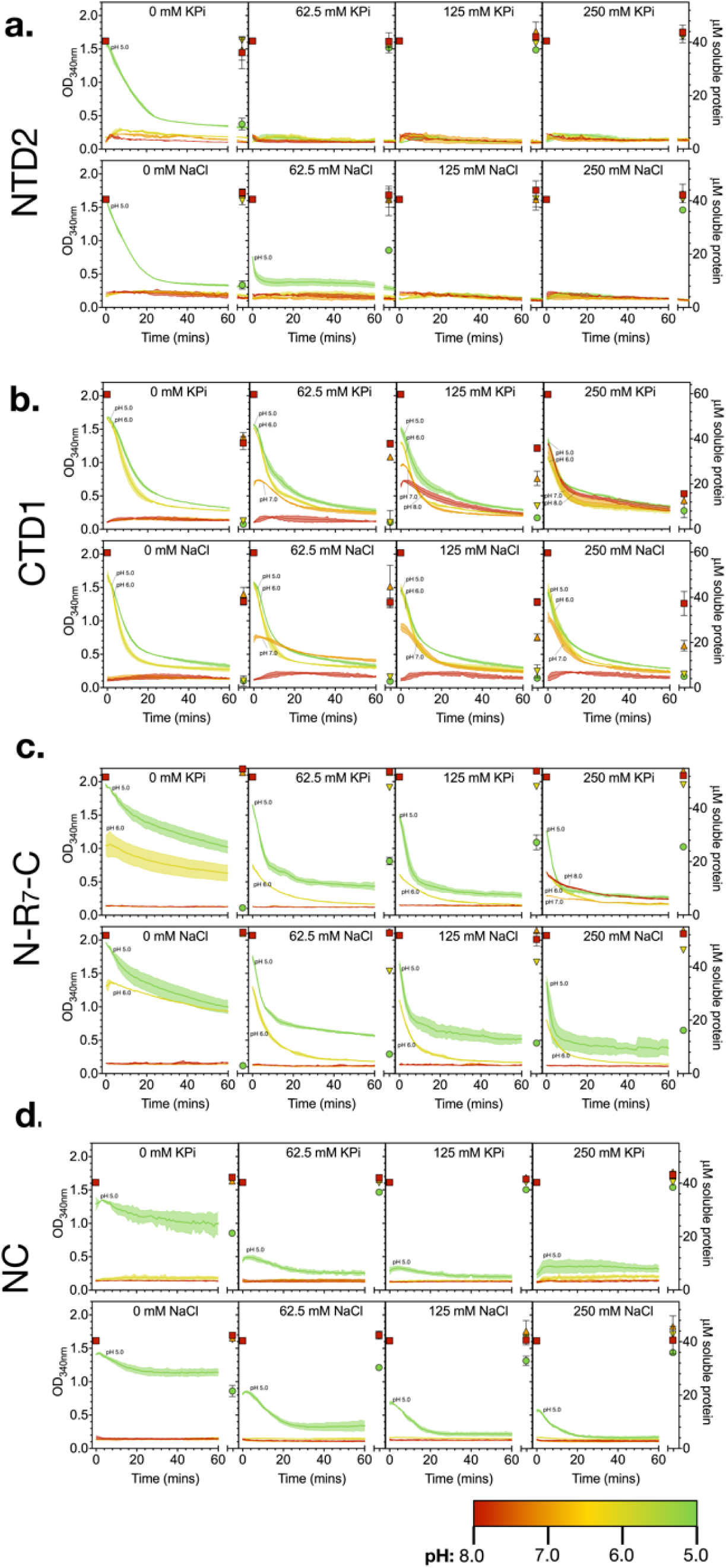
NTD2 (A), CTD1 (B), N-R_7_-C (C), and NC (D) characterization. The effect of pH and salt. Turbidity as measured by OD_340nm_ (left axis) over time at various pH values and salt concentrations. pH is indicated by colour: pH 5 (green), pH 6 (yellow), pH 7 (orange), pH 8 (red), as also indicated in the legend. The right axis shows protein concentration, as determined by nanodrop at OD_280nm_ before and after the assay, shown by pH 8: squares, pH 7: upwards triangles, pH 6: downwards triangles and pH 5: circles. Error bars show the standard deviation of three replicates in both cases.

NTD2 aggregated in the absence of NaCl or KPi at pH 5.0 (**Figure 5A**). Dimerization of spidroin N-terminal domains is induced via a protonation-induced dipole interaction ^35^. In the absence of salt, our results suggest that the formation of this dipole at pH 5.0 results in the undesirable aggregation of NTD2 domains, rather than simply dimerization. Stabilization of the NTD dipole at pH 5.0, through the addition of NaCl or KPi, appears to prevent this aggregation (**Figure 5)**. Indeed, electrolytes have been proposed to be important in stabilising local clusters of negative and positive charge on the NTD surface ^35^, with the addition of NaCl shifting the pKa for dimerization towards a more acidic pH ^36^.

Such aggregation of NTD2 likely also occurs in the context of a complete mini-spidroin. Indeed, the rate of assembly of N-R_7_-C into larger aggregates, indicated by decreasing turbidity, was substantially slower in the absence of NaCl or KPi (**Figure 5C**). A similar response was observed for a construct featuring both terminal domains, but no repetitive region (NC, **Figure 5D**). In contrast, CTD1 alone did not behave in this way (**Figure 5B**), suggesting this response is due to the effects of NTD2, which aggregated in this condition (**Figure 5A**).

N-R_7_-C also displays a large negative shift in thermal stability in the absence of salt at pH 5.0 (**Figure 6**), which is likely caused by the un-stabilised dipole at the N-terminal domain at this pH. Similar DSF experiments of NC, NTD2 and CTD1 appear to support this assessment (Supplementary Material **Figure S8**). Interestingly, we also observed a decrease in the TM of N-R_7_-C with increasing KPi, and to a much lesser extent higher concentrations of NaCl, at pH 8.0 and 7.0, suggesting the mini-spidroin is becoming destabilised by the addition of these salts at neutral pH (**Figure 6**). This effect is not observed at lower pH values which may be due to a stronger response to pH overall.

**Figure 6.**
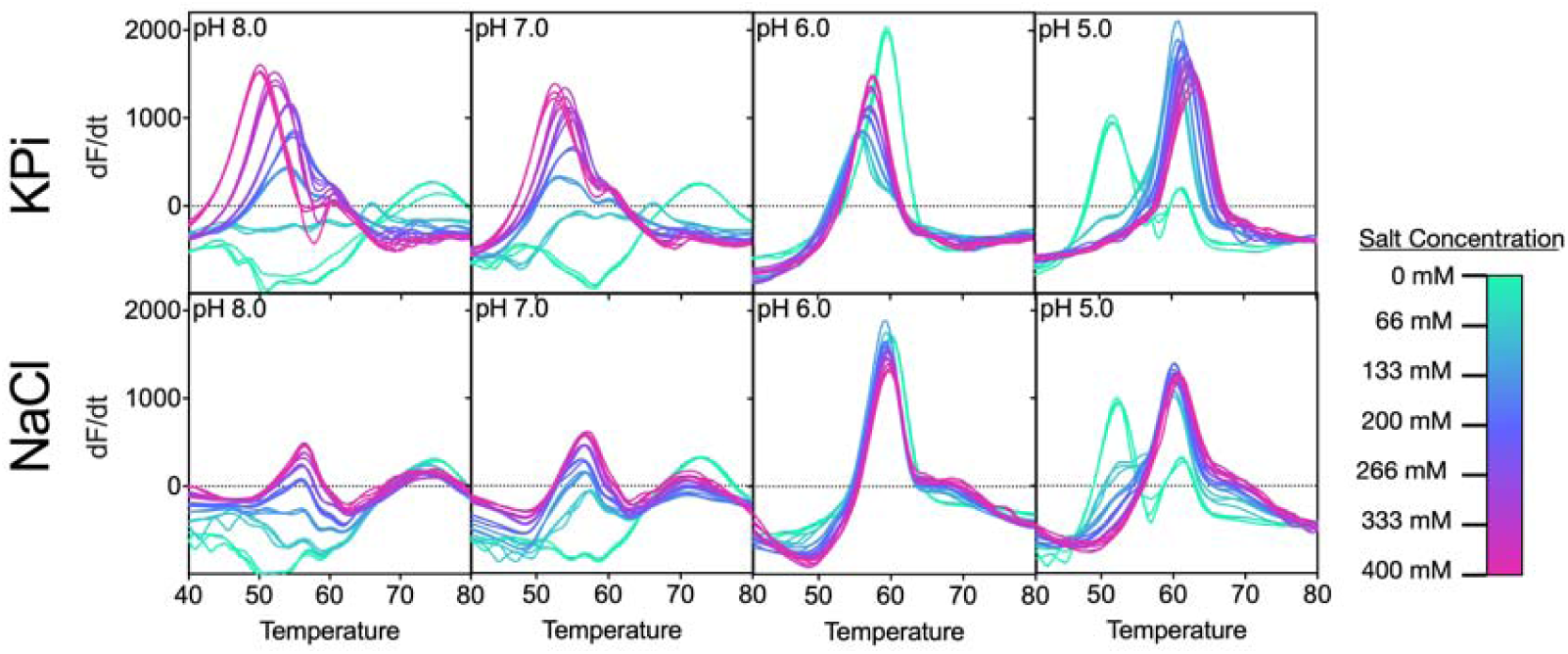
Differential scanning fluorimetry of N-R7-C at different pH values and NaCl or KPi concentrations. NaCl or KPi concentrations are indicated according to the colour chart. A large shift in the TM is observed at pH 5.0 in the absence of either NaCl or KPi, as indicated by the red arrow. Temperature is shown in °C. Negative signals in the DSF assay indicate a decreasing fluorescence signal (as dF/dT is plotted). This may be due exposed hydrophobic regions on the protein (either due to the native conformation of the protein or due to a fraction of the sample being denatured at the start of the assay) to which sypro orange can immediately bind, gradually releasing the dye as temperature increases, resulting in a loss of fluorescence signal.

Our results suggest the slow rate of assembly of N-R_7_-C into larger aggregates at pH 5.0 in the absence of NaCl or KPi is due to incorrect aggregation of the NTD2 domain, as observed with this domain alone, which likely inhibits the correct assembly of the mini-spidroin. Indeed, it has been indicated that the presence of NaCl in the spiders silk gland is important in preventing undesired aggregation ^37^.

A previous study has shown another NTD to behave in a similar way, with turbidity measurements of NTD much higher at pH 6 in the absence of salt, than either pH 7 or 6 in the presence of salt ^38^. The inclusion of this NTD in a mini-spidroin resulted in macroscopic structures forming earlier in a self-assembly assay in the absence of salt, but no faster in the presence of salt. In light of our results, possibly the earlier formation of macroscopic structures in the absence of salt could be due to fast, non-specific aggregation at the NTD, which is undesirable in a spinning process.

### 3.5 Both a pH drop and salting out are required for effective biomimetic wet spinning

Exchange of sodium and chloride ions for potassium and phosphate ions in the natural spinning process is proposed to induce salting out of the spidroins ^12–14^. To further examine the effect of increasing potassium phosphate concentration during biomimetic wet spinning, N-R_7_-C at 100 mg/mL was extruded via a needle into a range of buffer conditions at both pH 5.0 and pH 8.0, and the resulting fibers observed (**Figure 7**). Again, potassium phosphate was prepared at the relevant pH by mixing solutions of KH_2_PO_4_ and K_2_HPO_4_.

**Figure 7.**
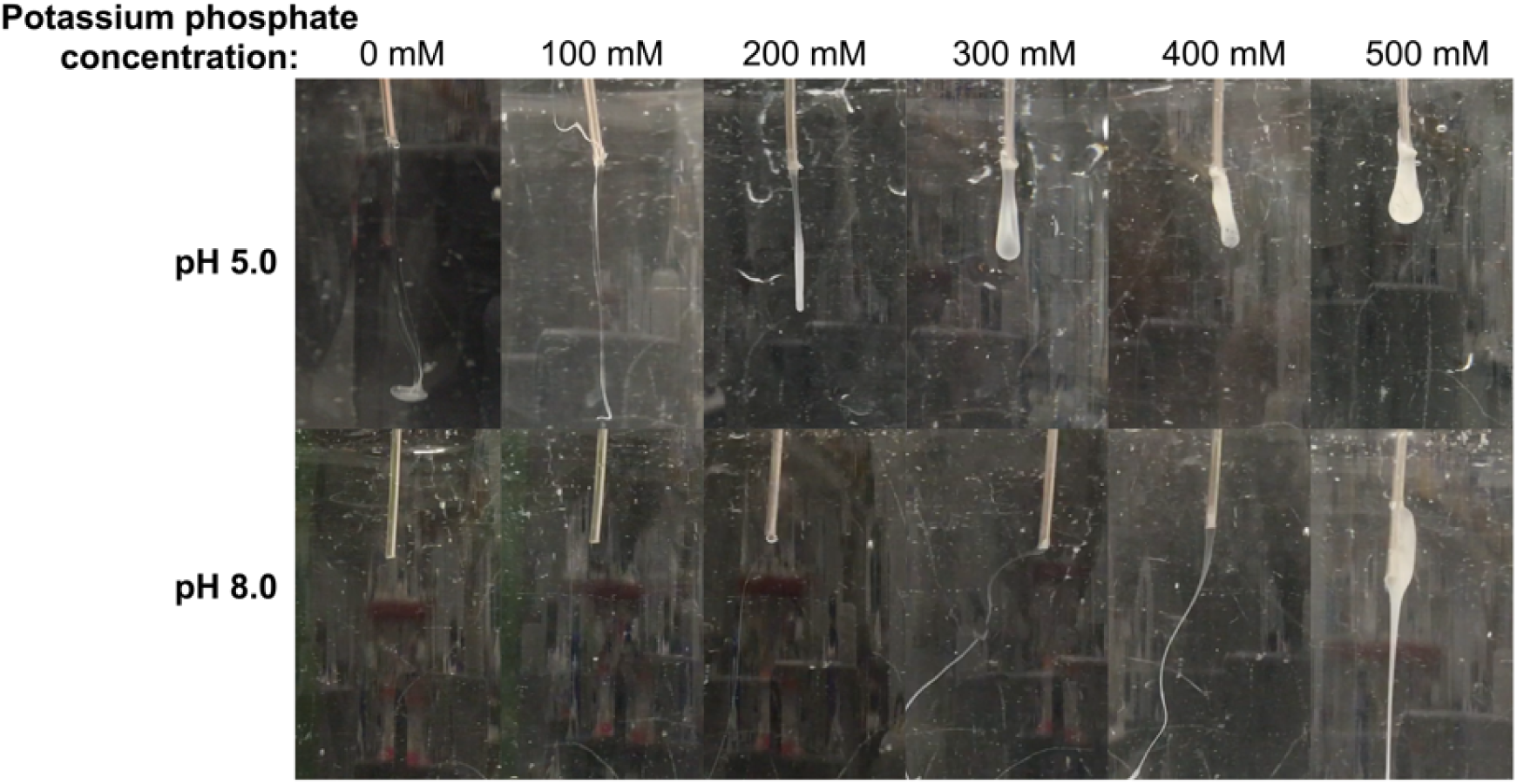
Wet spinning of N-R_7_-C. Different ranges of potassium phosphate concentrations were used at pH 5 and pH 8. 100 mg/mL (10 % w/v) to extrude N-R_7_-C at 25 µL/min through a blunted 16G needle. 50 mM sodium acetate or TrisHCl was used to at pH 5 and pH 8 respectively. Potassium phosphate was prepared at the relevant pH by mixing solutions of KH_2_PO_4_ and K_2_HPO_4_ in each case.

At pH 8.0, high (>300 mM) potassium phosphate concentrations resulted in visible aggregates that appeared to extrude from the needle as a fiber, likely due to salting-out of the mini-spidroin. The salting-out of N-R7-C appears similar to as observed for CTD1 alone at pH 8.0 and high potassium phosphate in the turbidity assays (**Figure 5**). However, in the absence of polymerisation of the mini-spidroin via NTD2, robust fibers were not formed resulting in only loosely associated aggregates. Collection of fibers at pH 8.0 at any KPi concentration was not possible, as they completely disintegrated upon contact in solution.

In contrast, at pH 5.0 with the addition of increasing concentrations of potassium phosphate, progressively more robust fibers formed. This resulted in an aggregated mass forming, rather than fibers, at higher potassium phosphate concentrations (>300 mM). In the absence of potassium phosphate (0 mM), a colloidal suspension was observed, likely a result of aggregation via NTD2 in this condition. Fibers could be collected or pulled and stretched from aggregates at the needle tip using tweezers (pH 5.0, >300 mM KPi only). Based on these observations, we conclude that both a drop to pH 5.0 in combination with salting-out is necessary for the formation of robust fibers which can be collected, and either of these triggers alone is insufficient. Importantly, these conditions allow an artificial dope with suitable rheological properties to be formed, which can then be mechanically drawn into a fiber.

We also note that a requirement for salting out has been shown not to be necessary for the self-assembly of some native and recombinant spidroins ^37–39^. Importantly, self-assembly is able to occur on a time-scale of hours, while fiber formation in a spinning process must occur over seconds, resulting in different requirements.

### 3.6 Biomimetic wet spinning of a complete mini-spidroin results in synthetic spider silk fibers

The production of fibers from N-R_7_-C by the extrusion into a biomimetic coagulation bath was tested. A coagulation bath consisting of 50 mM sodium acetate, 500 mM potassium phosphate pH 5.0 was used, inducing a pH drop in combination with salting-out. An aggregated mass formed at the tip of needle from which a fiber could be pulled by tweezers and collected continuously onto a rotating collector (**Supplementary Material 2-movie**). This is analogous to how spiders spin silk, with the silk pulled from the spinneret rather than pushed, in a process referred to as pultrusion ^40^. This use of mechanical force is also important in achieving strong, aligned fibers, evidenced by the need for post-spin drawing in many examples of synthetic spider silk spinning ^41^. The diameters of the fibers, as determined by light microscopy, varied between 14 and 51 μm. Engineering stress and strain of as-spun fibers were determined by tensile testing, with a mean ultimate tensile strength of 40.3 MPa, and a maximum of 78 MPa (**Figure 8**). The fibers showed a lower Young’s modulus than native spider silk (11.6 ± 0.7 GPa ^42^). Thinner fibers correlated with higher ultimate tensile strength (UTS), and higher values for Young’s modulus (R = −0.62 and R = −0.53 respectively, Supplementary Material **Figure S9 and S10**). A comparison with the mechanical properties obtained in other studies is shown in Supplementary Material **Table S1**. Scanning electron microscopy (SEM) of some fibers evidenced many aligned fibrils (**Figure 9A, 9B**). However, a second batch of fibers spun separately and imaged at higher resolution did not show this (**Figure 9C**). The presence of many aligned fibrils constituting the fiber would be promising as natural spider silk is thought to consist of a hierarchical structure with many silk fibrils, covered by a skin layer, making up a silk fiber ^43^.

**Figure 8.**
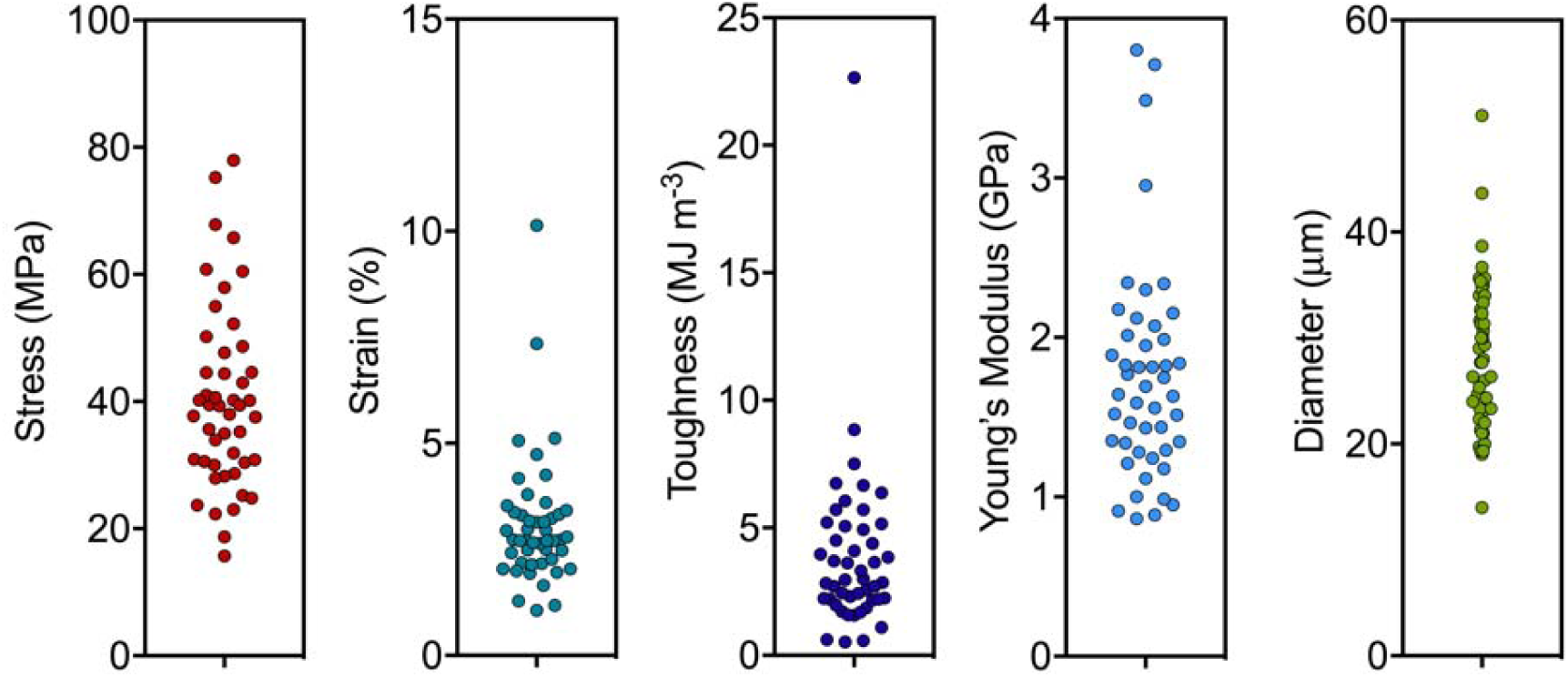
Mechanical testing of synthetic spider silk fibers from N-R_7_-C. Mechanical properties were calculated from stress strain curves (**Figure S9**).

**Figure 9.**
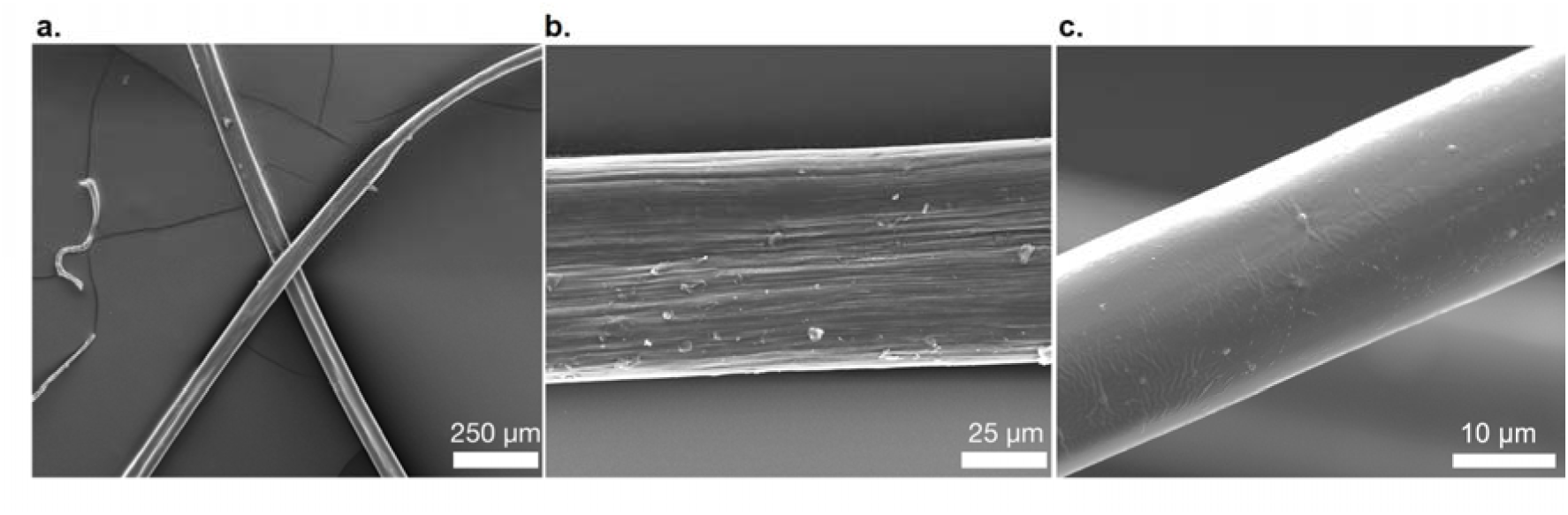
SEM images of spun fibers of N-R_7_-C. Scale bars show 250 μM (A), 25 μm (B) 10 μm (C). Aligned fibrils are observed at high magnification in some cases (B), but not others (C). The fiber shown in panel C represents a second batch of fibers, spun on a separate occasion to the fibers shown in A and B.

Wide-angle X-ray diffraction (WAXD) was conducted on a bundle of fibers to probe their crystallinity (Supplementary Material **Figure S11**). A broad amorphous region was observed along with two distinguishable diffraction peaks at approximately 10.4 and 22.6 degrees, indexed respectively as the (100) and (120) Bragg reflections of β-sheet crystallites as reported by Du et al ^44^. The average β-sheet crystallite size was calculated as 4.8 nm × 2.0 nm by application of the Scherrer equation on the deconvoluted peaks;

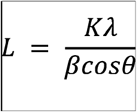

where L is the mean size of the crystallite domains, λ is the X-ray wavelength (1.542 Å), K is the dimensionless shape factor (taken as 0.9), β is the peak full width at half maximum (FWHM) and θ is the diffraction angle. The data were broadly in agreement with WAXD patterns of natural spider silk reported by Du et al ^44^.

Our results are in good agreement with previous work, which demonstrated the wet-spinning of a mini-spidroin with both terminal domains via a pH drop. We selected a pH drop to 5.0 for comparison with this work, and have more thoroughly investigated the effects of ionic strength on this spinning process, demonstrating this to be an important factor in this approach ^5^. The development of a spinning process utilising multiple smaller pH drops, or a continuous gradient could offer further improvements ^3^. In the development of such an approach it would likely be insightful to further consider the isoelectric point (pI) of the mini-spidroins and their constituent domains ^37^.

## 4. Conclusions

Biomimetic spinning offers a number of advantages over spinning using denaturing conditions, which arguably produces aggregates of denatured protein rather than correctly assembled spider silk. However, in order to be effective, we must understand the impact of pH, ion exchange and shear stress on biomimetic spinning dopes consisting of “complete” spidroins featuring both terminal domains. This work examines the production of a range of such mini-spidroins, and thoroughly investigates the conditions necessary for their biomimetic spinning. Similarities between the rheology of a mini-spidroin in this study, and that of native spider silk spinning dopes, suggest the inclusion of terminal domains is crucial in mimicking the spiders use of shear in the spinning duct to help achieve aligned fibrillar fibers. The terminal domains also allow polymerisation of a spidroin upon a pH drop. However, our results suggest that a pH drop alone is insufficient and a combination with salting-out, for which spiders use the exchange of sodium and chloride ions for potassium and phosphate ions, is critical for the production of robust fibers in a biomimetic wet spinning process. Using these conditions, biomimetic wet spinning allowed fibers to be spun continuously from a small mini-spidroin, with fibers formed from many aligned fibrils. However, fibers with diameters larger than native spider silk, and relatively poor mechanical properties, suggest an improved biomimetic spinning process is required. Our results provide the basis for the development of such a biomimetic spinning technique, which might combine shear with a biomimetic assembly buffer, as characterized here. Such a technique could offer substantial improvements to the quality of fibers achievable from small mini-spidroins, which are attractive for production at industrial scale since they can be produced at high yields. Future investigations into the underlying mechanisms by which ionic strength and pH drop facilitate fiber formation, could offer further improvements.

## Acknowledgments

WF and ADR acknowledge funding from the Defence, Science and Technology Laboratory (DSTL, UK Ministry of Defence, DSTLX1000101893). This is a contribution from the Manchester Centre for Synthetic Biology of Fine and Speciality Chemicals (SYNBIOCHEM) and acknowledges the Biotechnology and Biological Sciences Research Council (BBSRC) and Engineering and Physical Sciences Research Council (EPSRC) for financial support (Grant No. William). JJB, ADR, ET, RB and NSS acknowledge funding from DSTL, UK Ministry of Defence, project no. CDE100640.

## Author Contributions

NSS, RB, JJB and ET conceived the initial study design and supervised all aspects of the work. WF and ADR performed the experimental work. WF and ADR drafted the manuscript. NSS, RB, JJB and ET revised the manuscript. All authors reviewed and approved the final manuscript.

## Additional Information

### Conflicts of Interest

The authors declare no conflict of interest.

