## Supplementary material 1 for "The effect of terminal globular domains on the response of recombinant mini-spidroins to fiber spinning triggers"

### SUPPLEMENTARY FIGURES

**Figure S1 – NTD and CTD expression and purification**

Samples of the soluble fraction (Sol), insoluble fraction (Insol), flow-through (FT) and elution (E) following nickel affinity chromatography of E. coli cell lysate, after expression NTD1, NTD2, CTD1 and CTD2 domains analysed by SDS-PAGE. Over-expression of the proteins is indicated at the expected molecular weight by the red arrows in each case. All proteins were successfully purified via an N-terminal 6xHis tag, as shown in the lanes labelled P.

A. Expression and purification of NTD1, NTD2, CTD1 and CTD2. Initially part of the spidroin repetitive sequence was mistakenly cloned with the NTDs resulting in larger domains. NTD1 shows proteolysis in the soluble fraction and following purification.
B. Expression of NTD1 and NTD2 with the excess repetitive sequence removed, resulting in a smaller purified domain. The smaller version of NTD2 was used in subsequent work. While NTD1 no longer shows proteolysis, protein aggregates were observed following purification. For this reason NTD2 was chosen for characterisation and for use in mini-spidroins.

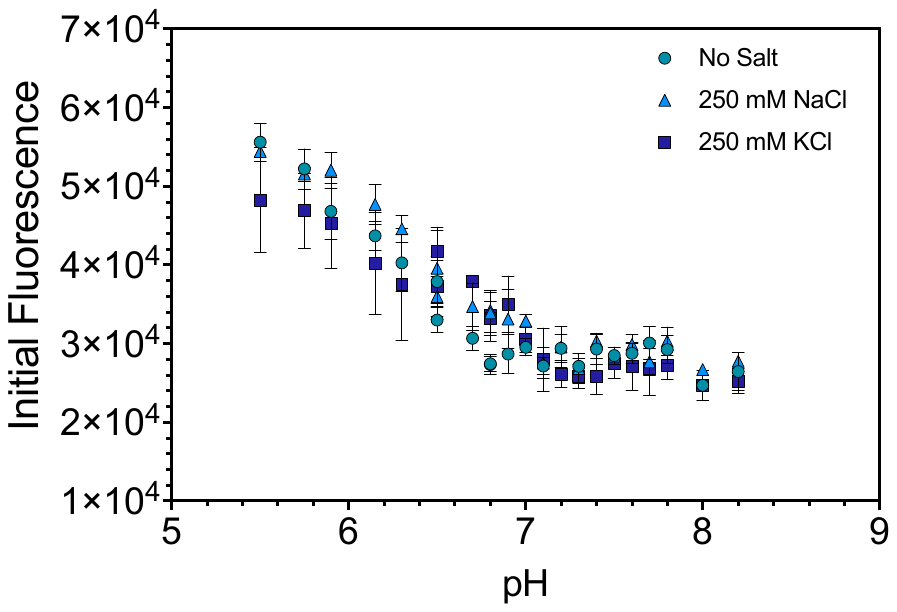

**Figure S2** **– Initial fluorescence of CTD1 in DSF assay.**

Sypro orange fluorescence upon binding to hydrophobic regions of proteins. A higher initial fluorescence at lower pH suggests more exposed hydrophobic regions of CTD1 at those pH’s. The assay was performed in the presence of 250 mM NaCl and KPO_4_, and without salt as indicated.

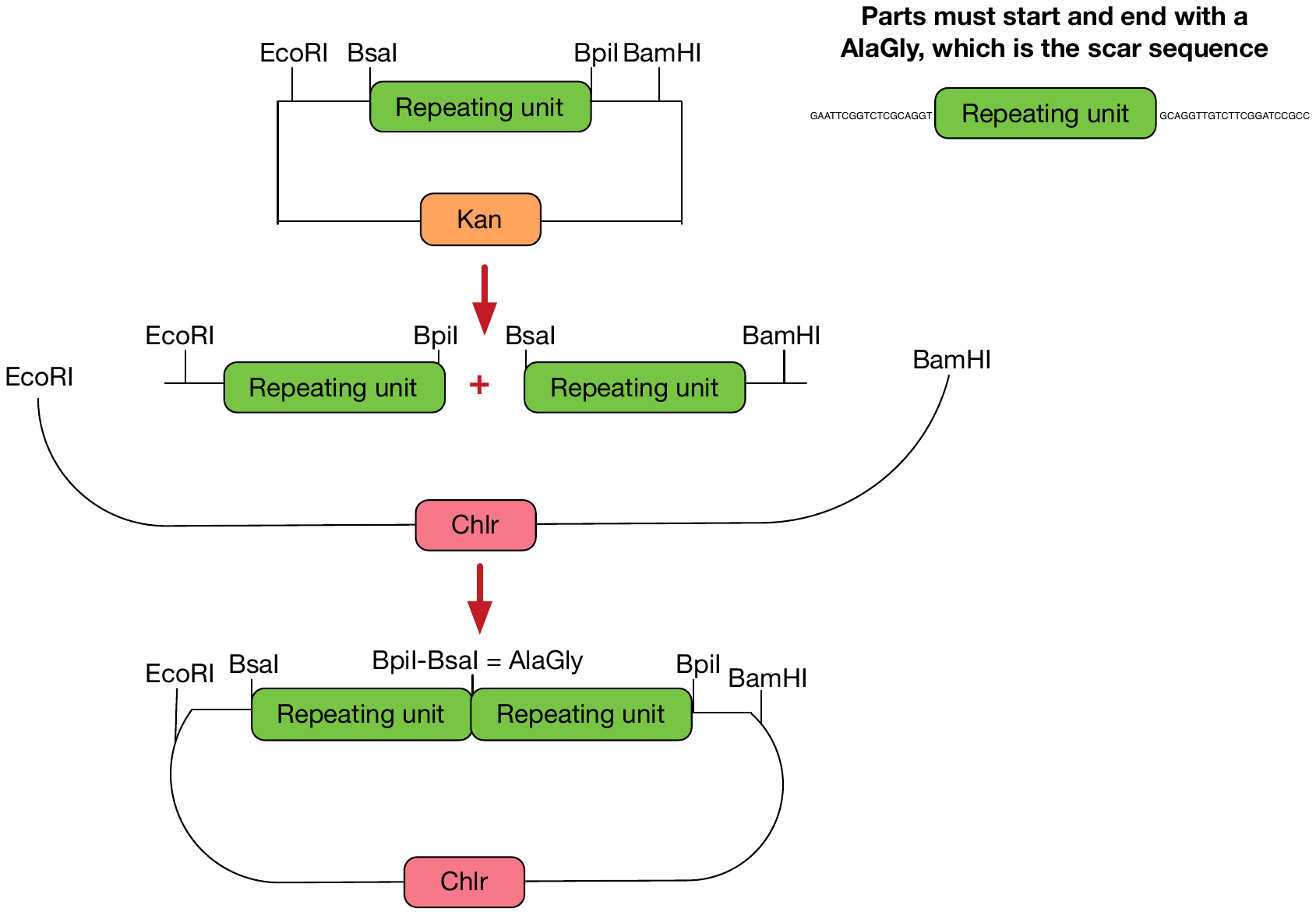

**Figure S3 – TypeIIS cloning scheme for pseudo-scarless duplication of repetitive sequences**

Scheme shows the use of restriction enzymes which can be used to carry out pseudo-scarless duplication of repetitive regions. This is achieved by designing the cut sites of the typeIIS restriction enzymes (BsaI and BpiI), to cut at a sequence coding for an alanine and a glycine, which will be present at the position regardless.

**Figure S4 – pTE1253 (NTD2-BsaI-CTD1-pNIC28) mini-spidroin expression vector.**

A. Plasmid map of pTE1253. A dashed box indicates the relevant section for mini-spidroin expression, shown in B.
B. Region for cloning repetitive sections into the BsaI sites, for the construction of mini-spidroins with both terminals.

**Figure S5 – Expression of mini-spidroin constructs after four hours expression.** SDS-PAGE of soluble (S) and insoluble (I) fractions of *E. coli* lysate following expression of various mini-spidroins for four hours at 20 ^o^C. Overexpressed proteins at the expected molecular weight are indicted by red arrows. Where no protein is detected the expected size is indicated by the red boxes.

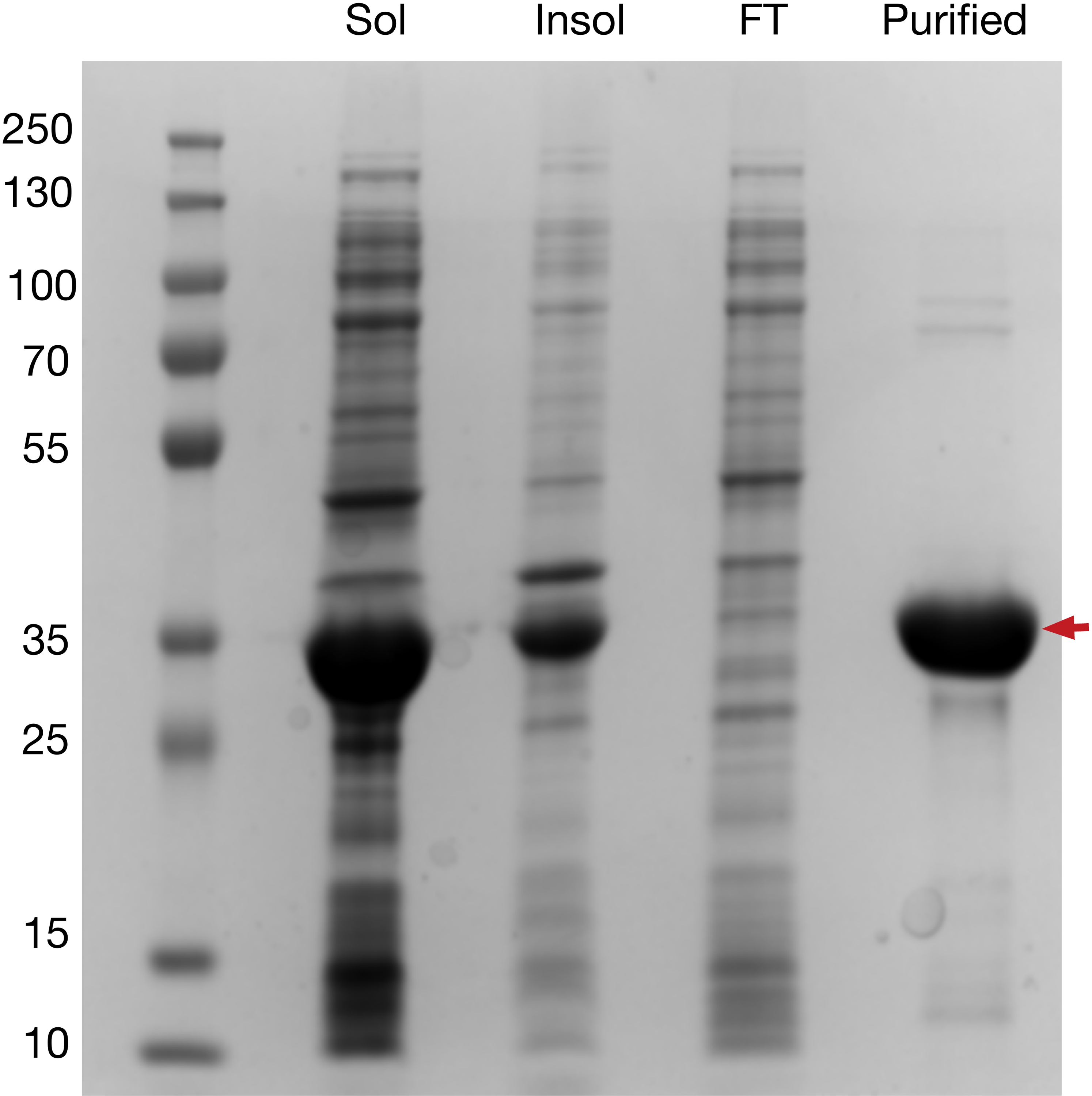

**Figure S6 – Ni IMAC purification of N-R_7_-C mini-spidroin.**

Samples of the soluble fraction (Sol), insoluble fraction (Insol), flow-through (FT) and elution (Purified) following nickel affinity chromatography of *E. coli* cell lysate, after expression of the mini-spidroin.

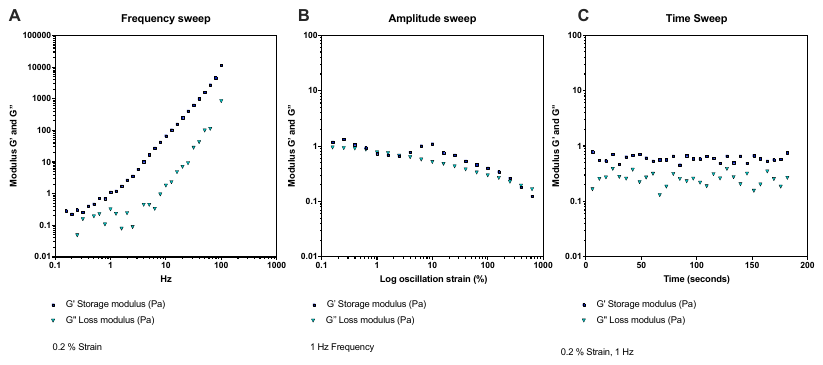

**Figure S7 – Preliminary frequency, amplitude and time sweeps of N-R_7_-C at 100 mg/mL.** Fixed parameters are displayed below each figure. The storage modulus (G’), and loss modulus (G’’) was recorded in each case. **A.** Frequency sweep suggests that the sample behaves like a weak gel, as G’ is greater than G’’. Different effects are observed at slow and fast time scales. **B.** Amplitude sweep also suggests the sample behaves as a weak gel, as G’ is greater than G’’. The weak gel appears to bear at approximately 400 % strain. **C.** Time sweep was conducted directly after the amplitude sweep with the same sample, and shows no memory of the gel breaking.

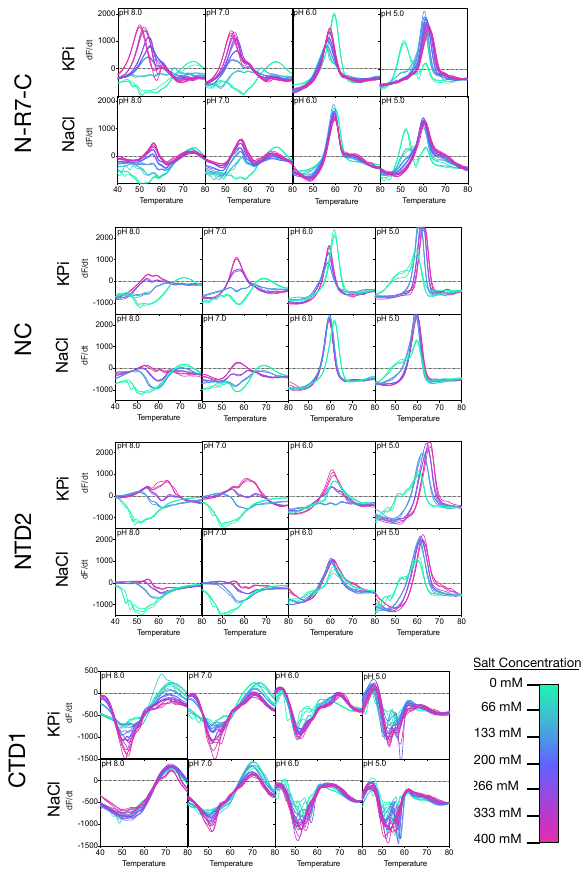

**Figure S8– DSF of NTD2, CTD1, NC and N-R_7_-C at different pH’s and salt concentrations** Three replicates are plotted for each condition. Temperature is shown in ^o^C. NaCl or KPi concentrations are indicated according to the colour chart. Negative signals in the DSF assay indicate a decreasing fluorescence signal (as dF/dT is plotted). This may be due exposed hydrophobic regions on the protein (either due to the native conformation of the protein or due to a fraction of the sample being denatured at the start of the assay) to which sypro orange can immediately bind, gradually releasing the dye as temperature increases, resulting in a loss of fluorescence signal. A shift in TM is observed at pH 5.0 for N-R7-C, NC or NTD2 in the absence of NaCl or KPO_4_, visible either as a distinct peak or as a shoulder towards a lower TM. This shift is not visible for CTD1, suggesting the response is caused by the presence of NTD2 in N-R7-C and NC.

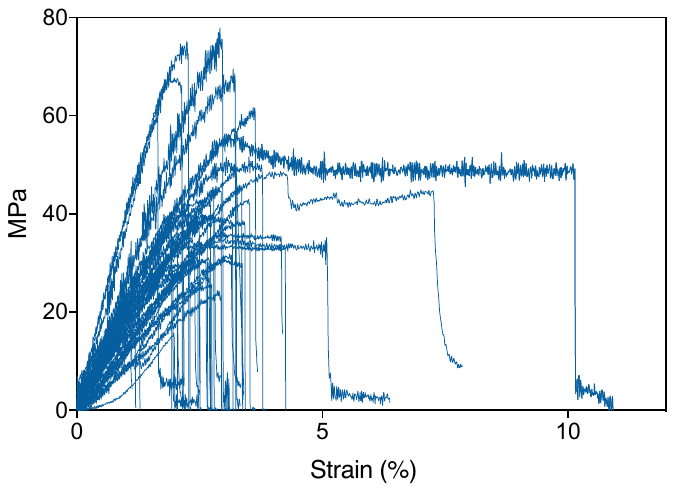

**Figure S9 – Stress strain curves for tensile tests of N-R_7_-C fibers.** All experiments carried out on the same day with environmental conditions of 28 ^o^C, 52 % humidity. Fibers were mounted on cardboard frames and mounted into a tensile testing machine (Instron 3344; Instron Ltd.), equipped with a 10 N load cell. Tensile tests were performed at a rate of 0.5 mm/min. Mechanical properties calculated using diameters estimated by light microscopy.

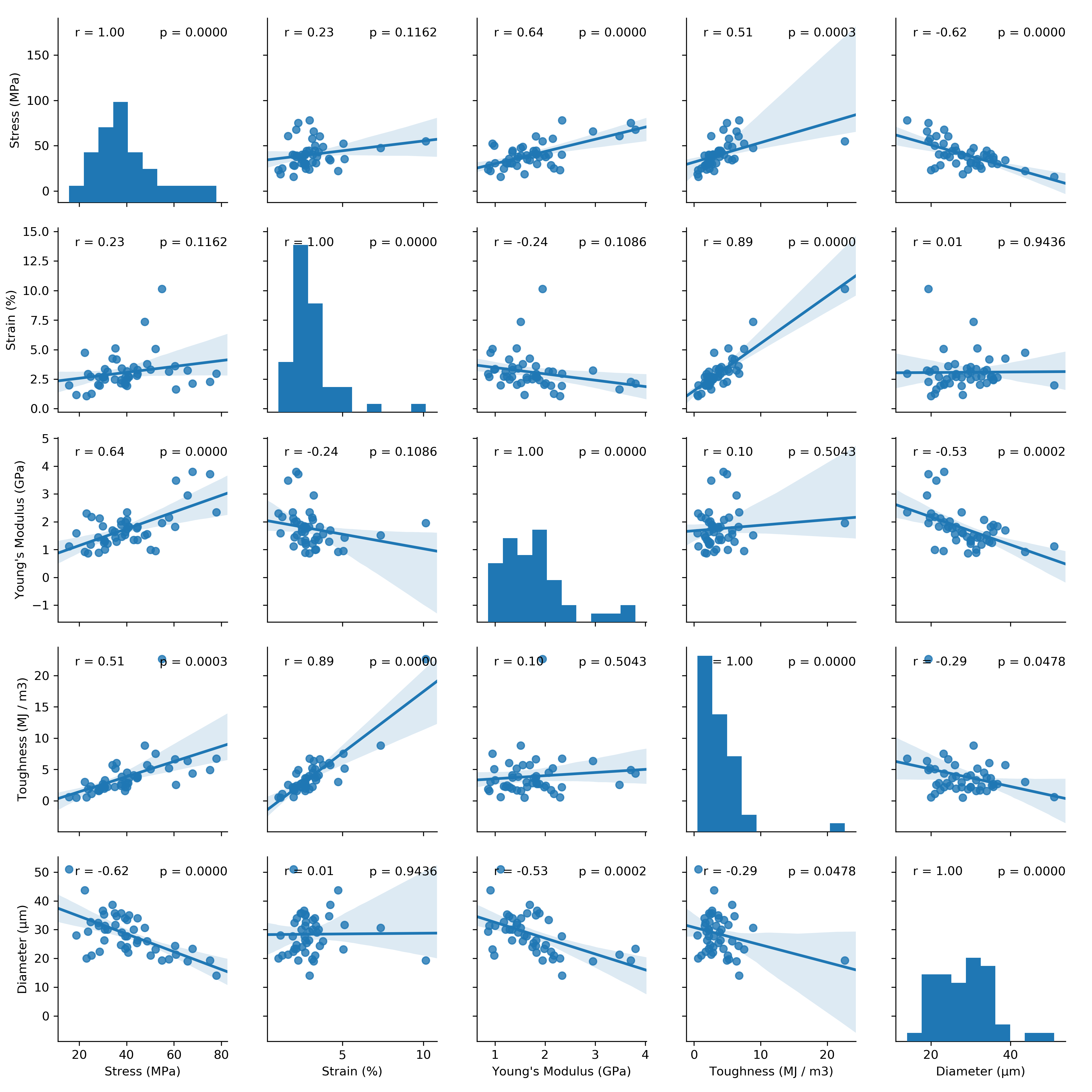

**Figure S10 – Correlation matrix of mechanical properties for fibers spun from N-R7-C**. Generated using the python package Seaborn. Correlation coefficient (r), and significance level (p) for each pair of factors is shown. Histograms showing the distribution of each factor are shown on the diagonal.

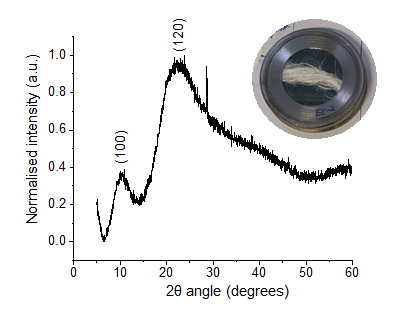

**Figure S11.** Wide-angle X-ray diffraction (WAXD) pattern of a bundle of fibers spun from spun from N-R7-C. **Inset:** fiber bundle on sample holder

|  | **N-R7-C (this study)** | **Native Dragline silk A. trifasciata** | **NT2RepCT** | **(AQ)24** | **N1L(AQ)24NR3** | **Synthetic 96-mer** | **Synthetic 192-mer** |
| --- | --- | --- | --- | --- | --- | --- | --- |
| **Reference** | This study | ^1^ | ^2^ | ^3^ | ^3^ | ^4^ | ^4^ |
| **Diameter (um)** | 28 ± 7 | ~3 | 12 ± 2 | 22 ± 2 | 20 ± 6 | 6.3 ± 0.7 | 5.7 ± 1.3 |
| **Extensibility (%)** | 3 ± 1.5 | 17 ± 0.04 | 37 ± 5 | 65 ± 6 | 54 ± 15 | 23 ± 7 | 22 ± 8 |
| **Strength (MPa)** | 40 ± 14 | 890 ± 130 | 162 ± 8 | 280 ± 48 | 308 ± 131 | 525 ± 83 | 1031 ± 111 |
| **Toughness (MJ m-3)** | 3.9 ± 3.4 | 100 ± 40 | 45 ± 7 | 110 ± 24 | 90 ± 29 | 91 ± 30 | 114 ± 58 |
| **Young's modulus (GPa)** | 1.8 ± 0.7 | 11.6 ± 0.7 | 6 ± 0.8 | 0.4 ± 0.1 | 5 ± 2 | 7.8 ± 1.3 | 13.7 ± 3 |

Supplementary Table 1 – A comparison of the mechanical properties obtained in this study, with an example of native spider silk and other recombinant spider silk fibers.

>NTD2_ unstructured_repetitive_region

MHHHHHHSSGVDLGTENLYFQSMALGQANTPWSSKENADAFIGAFMNAASQSGAFSSDQIDDMSVISNTLMAAMDNMGGRITQSKLQALDMAFASSVAEIAVADGQNVGAATNAISDALRSAFYQTTGVVNNQFITGISSLIGMFAQVSGNEVSYSSAGSSSAAASEAVSAGQGPAAQPVYAPSGASAAAAAASGAAPAIQQAYERGGSGSAAAAA*

>NTD2

MHHHHHHSSGVDLGTENLYFQSMALGQANTPWSSKENADAFIGAFMNAASQSGAFSSDQIDDMSVISNTLMAAMDNMGGRITQSKLQALDMAFASSVAEIAVADGQNVGAATNAISDALRSAFYQTTGVVNNQFITGISSLIGMFAQVSGNEV*

>CTD1

MHHHHHHSSGVDLGTENLYFQSMGSGPGQIYYGPQSVAAPAAAAASALAAPATSARISSHASALLSNGPTNPASISNVISNAVSQISSSNPGASACDVLVQALLELVTALLTIIGSSNIGSVNYDSSGQYAQVVTQSVQNAFA*

>NC

MHHHHHHSSGVDLGTENLYFQSMALGQANTPWSSKENADAFIGAFMNAASQSGAFSSDQIDDMSVISNTLMAAMDNMGGRITQSKLQALDMAFASSVAEIAVADGQNVGAATNAISDALRSAFYQTTGVVNNQFITGISSLIGMFAQVSGNEVGSGAGGGSGPGQIYYGPQSVAAPAAAAASALAAPATSARISSHASALLSNGPTNPASISNVISNAVSQISSSNPGASACDVLVQALLELVTALLTIIGSSNIGSVNYDSSGQYAQVVTQSVQNAFAGS*

>N-R7-C

MHHHHHHSSGVDLGTENLYFQSMALGQANTPWSSKENADAFIGAFMNAASQSGAFSSDQIDDMSVISNTLMAAMDNMGGRITQSKLQALDMAFASSVAEIAVADGQNVGAATNAISDALRSAFYQTTGVVNNQFITGISSLIGMFAQVSGNEVAGGGAGQGGQGGYGRGGYGQGGAGQGGAGAAAAAAAAGGAGQGGQGGYGQGGYGQGGAGQGGAAAAAAAAAGGAGQGGYGRGGAGQGGAGGSGPGQIYYGPQSVAAPAAAAASALAAPATSARISSHASALLSNGPTNPASISNVISNAVSQISSSNPGASACDVLVQALLELVTALLTIIGSSNIGSVNYDSSGQYAQVVTQSVQNAFAGS*

>N-R10-C

MHHHHHHSSGVDLGTENLYFQSMALGQANTPWSSKENADAFIGAFMNAASQSGAFSSDQIDDMSVISNTLMAAMDNMGGRITQSKLQALDMAFASSVAEIAVADGQNVGAATNAISDALRSAFYQTTGVVNNQFITGISSLIGMFAQVSGNEVAGGGAGQGGQGGYGRGGYGQGGAGQGGAGAAAAAAAAGGAGQGGQGGYGQGGYGQGGAGQGGAAAAAAAAAGGAGQGGYGRGGAGQGGAAAAAAAAGAGQGGYGGQGAGQGGAGAAAAAAAAGGSGPGQIYYGPQSVAAPAAAAASALAAPATSARISSHASALLSNGPTNPASISNVISNAVSQISSSNPGASACDVLVQALLELVTALLTIIGSSNIGSVNYDSSGQYAQVVTQSVQNAFAGS*

>N-R13-C

MHHHHHHSSGVDLGTENLYFQSMALGQANTPWSSKENADAFIGAFMNAASQSGAFSSDQIDDMSVISNTLMAAMDNMGGRITQSKLQALDMAFASSVAEIAVADGQNVGAATNAISDALRSAFYQTTGVVNNQFITGISSLIGMFAQVSGNEVAGGGAGQGGYGRGGAGQGGAAAAGAGQGGYGGQGAGQGGAGAAAAAAAAGGAGQGGQGGYGRGGYGQGGAGQGGAGAAAAAAAAGGAGQGGQGGYGQGGYGQGGAGQGGAAAAAAAAAGGAGQGGYGRGGAGQGGAAAAAAAAGAGQGGYGGQGAGQGGAGAAAAAAAAGGSGPGQIYYGPQSVAAPAAAAASALAAPATSARISSHASALLSNGPTNPASISNVISNAVSQISSSNPGASACDVLVQALLELVTALLTIIGSSNIGSVNYDSSGQYAQVVTQSVQNAFAGS*

>N-R36-C

MHHHHHHSSGVDLGTENLYFQSMALGQANTPWSSKENADAFIGAFMNAASQSGAFSSDQIDDMSVISNTLMAAMDNMGGRITQSKLQALDMAFASSVAEIAVADGQNVGAATNAISDALRSAFYQTTGVVNNQFITGISSLIGMFAQVSGNEVAGGAGQGGQGGYGRGGYGQGGAGQGGAGAAAAAAAAGGAGQGGQGGYGQGGYGQGGAGQGGAAAAAAAAAGGAGQGGYGRGGAGQGGAAAAGAGQGGYGGQGAGQGGAGAAAAAAAAGGAGQGGQGGYGRGGYGQGGAGQGGAGAAAAAAAAGGAGQGGQGGYGQGGYGQGGAGQGGAAAAAAAAAGGAGQGGYGRGGAGQGGAAAAAAAAGAGQGGYGGQGAGQGGAGAAAAAAAAGGSGPGQIYYGPQSVAAPAAAAASALAAPATSARISSHASALLSNGPTNPASISNVISNAVSQISSSNPGASACDVLVQALLELVTALLTIIGSSNIGSVNYDSSGQYAQVVTQSVQNAFAGS*

Larger proteins are not shown, but available through the plasmid files in the supplementary information.

Supplementary Table 2 – Protein sequences for a selection of proteins used in this study. Complete plasmid files are also available as an additional supplementary file.
